# Experimental and computational mapping reveals preferential nanobody targeting of the intracellular cavity of the amino acid transporter BasC

**DOI:** 10.64898/2026.11.11.765408

**Authors:** Adrià Nicolàs-Aragó, Daniel Doherty, Patricia Gómez-Gejo, Paola Bartoccioni, Els Pardon, Jan Steyaert, Ekaitz Errasti-Murugarren, Manuel Palacín, Joana Fort

## Abstract

Nanobodies are valuable tools for stabilizing and investigating membrane transporters, but the systematic characterization of large nanobody collections remains experimentally demanding. Here, we combined biochemical and transport assays, experimental structural data and AlphaFold3 modelling to characterize a repertoire of 29 nanobodies targeting the bacterial amino acid transporter BasC. By exploiting the random orientation and asymmetric apparent substrate affinities of BasC in proteoliposomes, comparison of nanobody-mediated influx and efflux inhibition enabled experimental assignment of binding sidedness. The collection showed a marked bias towards inhibition from the intracellular face, whereas only two, Nb58 and Nb78, were identified as extracellular binders. Compatibility assays and a 6.1 Å crystallographic map of the BasC–Nb53–Nb58 complex supported the simultaneous binding of Nb53 and Nb58 to opposite sides of the transporter and validated the functional sidedness assignment. AlphaFold3 generated high-confidence interaction models for 16 of the 29 complexes. Ten of these models targeted the large intracellular cavity, revealing recurrent epitope clusters. The predicted binding modes of Nb53, Nb71 and Nb74 agreed with available experimental structures, supporting the accuracy of these predictions. Notably, these three nanobodies had previously provided the most informative structural and smFRET data. Our results show that transport-based sidedness measurements together with confidence-filtered structural predictions can define the functional and structural landscape of a nanobody repertoire. This integrated approach may provide a virtual structural screening strategy to prioritize nanobodies, identify dominant epitope regions and select potentially compatible combinations for experimental and structural studies.

## Introduction

Nanobodies are valuable tools for studying membrane proteins because they can recognize conformational epitopes, stabilize specific states and facilitate structural characterization (Pardon et al. 2014). For membrane transporters, they can also provide functional probes linking substrate transport to conformational changes. However, identifying the most informative binders within a nanobody repertoire requires more than measuring affinity: their binding location, effects on transport and compatibility with other nanobodies can also be established.

Experimental discovery platforms have expanded access to nanobodies against membrane proteins, including mRNA/cDNA display applied to GLUT1 (Doshi et al. 2014), synthetic single-domain antibody (sybody) selection against SLC and ABC transporters (Zimmermann et al. 2018), and yeast surface display for conformationally selective binders to GPCRs (McMahon et al. 2018). These platforms expand access to membrane-protein binders, but systematic characterization of the resulting repertoires remains experimentally demanding.

The bacterial amino acid exchanger BasC provides an experimentally accessible model for investigating L-type amino acid transporter mechanisms. Previous structural and functional studies have established its asymmetric substrate interactions and identified key determinants of cytoplasmic gating (Errasti-Murugarren et al. 2019; Fort et al. 2025). Nanobodies contributed to these studies as structural chaperones and mechanistic probes. Nevertheless, the broader repertoire from which these binders were selected remains incompletely characterized, offering an opportunity to examine how nanobody diversity relates to binding location and functional effects.

Structure prediction offers a complementary approach to this experimental characterization. AlphaFold-based methods can recover antibody–antigen binding modes for a subset of complexes, and AlphaFold3 (AF3) has improved prediction accuracy for these interactions (Yin y Pierce 2024; Abramson et al. 2024). However, their variable performance makes comparison with experimental evidence essential. Here, we combine biochemical characterization, transport-based sidedness assays, nanobody compatibility experiments, low-resolution crystallography and AF3 modeling to investigate 29 BasC nanobodies. Our aim is to map their functional and structural diversity and assess whether confidence-filtered predictions can help prioritize nanobodies for mechanistic and structural studies.

## Results

### Generation and biochemical characterization of a BasC nanobody repertoire

To explore how nanobodies can probe and modulate LAT-family transporters, a panel of nanobodies against the bacterial LAT homologue BasC was generated (Errasti-Murugarren et al. 2019). Immunization was performed using purified BasC reconstituted into liposomes, thereby presenting the transporter in a membrane environment inserted randomly (Bartoccioni et al. 2019). Nanobody discovery yielded 73 ELISA-positive clones, which were sequence-analyzed and grouped into 27 families primarily according to CDR3 similarity.

Representative clones from these families were selected for further biochemical characterization, resulting in the panel of 29 nanobodies analyzed in this study (Supplementary Table S1) (Pardon et al. 2014).

CDR3 sequence comparison of the selected nanobody panel revealed a highly diverse repertoire (Supplementary Fig. S2; Supplementary Table S1). A few nanobodies showed clear sequence relatedness, including Nb50–Nb51, Nb61–Nb62 and Nb65–Nb68, whereas most of the remaining nanobodies displayed highly divergent CDR3 sequences and were broadly distributed across the dendrogram. This sequence diversity provided a useful framework for subsequently assessing whether CDR3 relatedness correlates with recognition of common or distinct BasC epitopes. Representative nanobodies were expressed and purified, and their interaction with BasC was analyzed using surface plasmon resonance (SPR) (Table 1 and Supplementary Table S3).

**Table 1.**
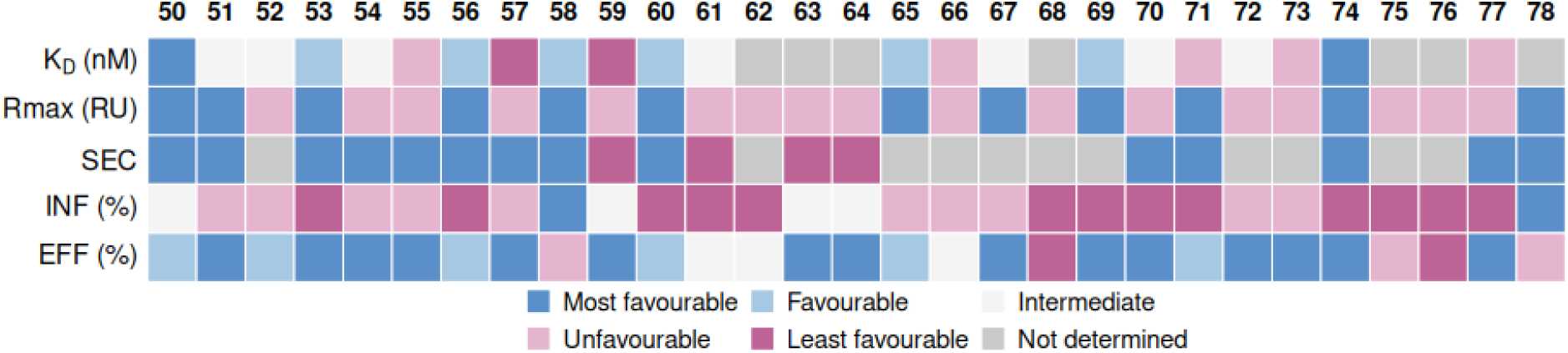
Summary of the biochemical and functional characterization of BasC nanobodies. The heatmap summarizes apparent equilibrium dissociation constants (K_D_) and maximum binding responses (R_max_) measured by surface plasmon resonance using detergent-solubilized BasC in the absence of L-alanine, stable complex formation assessed by size-exclusion chromatography (SEC), and residual influx (INF) and efflux (EFF) transport activities relative to the control without nanobody. Colors indicate parameter-specific categories, with lower K_D_, higher R_max_, stable complex formation and lower residual transport activity classified as more favorable. Color categories summarize the biochemical and functional profiles and do not indicate statistical significance. Grey cells indicate measurements that could not be determined. Quantitative values and additional experimental details are provided in Supplementary Table S3.

Binding of the nanobody panel to detergent-solubilized BasC was systematically evaluated by surface plasmon resonance (SPR) using a stepwise screening strategy. In an initial screen, all nanobodies were immobilized and tested at two BasC concentrations to identify the best binders and obtain a preliminary assessment of their binding behavior. Selected nanobodies were subsequently analyzed over an expanded concentration series to determine their kinetic parameters and equilibrium dissociation constants (KD) (Table 1). Representative sensorgrams for Nb53, Nb58 and Nb71 are shown in Supplementary Figure S4.

Together, these results establish a diverse repertoire of BasC-binding nanobodies with distinct kinetic profiles and complex formation behavior, providing the basis for subsequent functional and structural analyses.

### Assignment of BasC nanobody binding sidedness by transport and compatibility assays

To determine from which side of the membrane individual nanobodies affect BasC function, transport assays were performed using BasC reconstituted into proteoliposomes. BasC is incorporated into proteoliposomes with a random orientation and shows higher apparent substrate affinity at the extracellular side than at the intracellular side (Bartoccioni et al., 2019). Thus, influx and efflux measurements performed at low substrate concentrations can be used to selectively measure right-side-out and inside-out BasC molecules, respectively. This approach was previously validated using Nb74, whose binding to the intracellular side of BasC was established structurally, allowing the two transporter orientations in proteoliposomes to be distinguished (Errasti-Murugarren et al. 2019)

In the influx assay, 10 μM radiolabelled amino acid was added to the external medium, and uptake into proteoliposomes mediated by right-side-out BasC molecules was measured. In the efflux assay, proteoliposomes were preloaded with radiolabelled substrate, and release into the external medium mediated by inside-out BasC molecules was monitored (Figure 1 A). Nanobodies were added exclusively to the external medium. Therefore, nanobodies that inhibited influx were classified as extracellular binders (Figure 1B and Table 1), whereas those that inhibited efflux were classified as intracellular binders (Figure 1B and Table 1).

**Figure 1.**
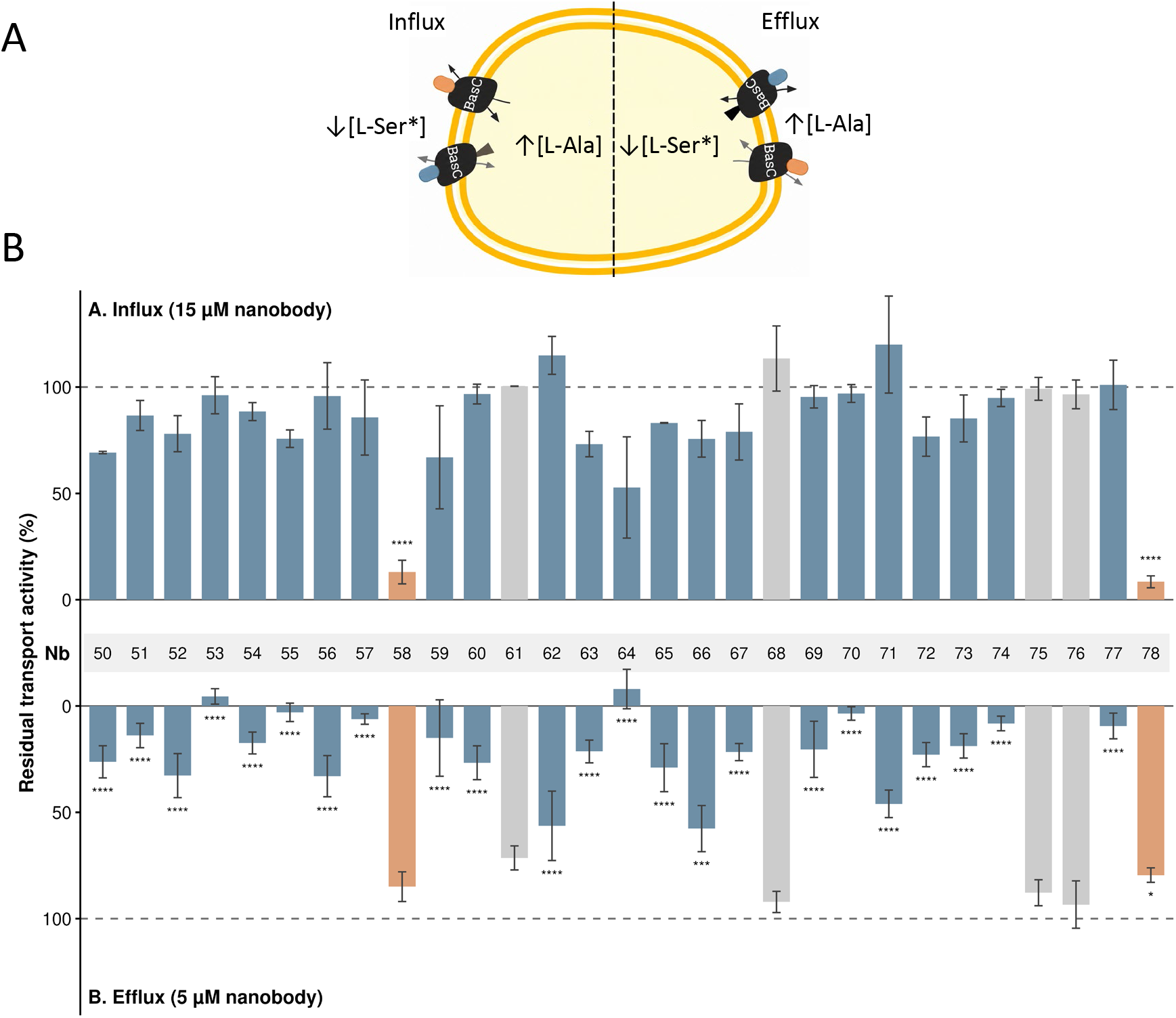
Functional characterization of BasC nanobodies by influx and efflux inhibition assays. **(A)** Experimental strategy for assessing nanobody-binding sidedness in randomly oriented BasC–GFP proteoliposomes. Influx was measured using 10 μM external L-[^3^H]-serine in proteoliposomes preloaded with 4 mM L-alanine. Efflux was measured from proteoliposomes preloaded with radiolabeled L-serine following addition of 4 mM external L-alanine. Under these conditions, influx and efflux preferentially report transport by right-side-out and inside-out BasC, respectively. Nanobodies were added exclusively to the extraliposomal medium. Orange and blue symbols illustrate nanobodies recognizing the extracellular and intracellular faces of BasC, respectively. **(B)** Residual influx and efflux activities following incubation with individual nanobodies at 15 μM and 5 μM, respectively. Activities were normalized to the corresponding control without nanobody (100%; dashed lines). Influx is plotted upwards and efflux downwards for visual comparison. Bars represent mean ± s.e.m. from 2–12 independent experiments for influx and 3–14 independent experiments for efflux, depending on the nanobody. Each condition was compared with the corresponding control without nanobody using Dunnett’s multiple comparisons test, separately for influx and efflux. Orange denotes nanobodies significantly inhibiting influx, including those also significantly inhibiting efflux; blue denotes nanobodies significantly inhibiting efflux only; and grey denotes those without significant inhibition in either assay. Asterisks indicate statistically significant differences from the corresponding control without nanobody: *P* < 0.05 (*), *P* < 0.001 (***) and *P* < 0.0001 (****).

Transport measurements revealed distinct effects of the nanobody repertoire on BasC activity. Nb58 and Nb78 strongly inhibited influx, supporting their assignment as extracellular binders, whereas most nanobodies inhibited efflux, consistent with intracellular binding. Nb59 and Nb64 significantly inhibited efflux, although they also showed a tendency to reduce influx activity. This tendency was further investigated in dose–response assays, which revealed partial influx inhibition, leaving their binding sidedness undetermined (Supplementary Figure S5). Four nanobodies (Nb61, Nb68, Nb75 and Nb76) did not significantly affect either transport assay under the conditions tested.

Influx dose–response assays showed that Nb58 and Nb78 produced near-complete inhibition, with IC_50_ values of 9.44 ± 2.42 and 34.8 ± 8.01 nM, respectively (Supplementary Figure S5). To further assess their extracellular assignment, nanobody compatibility was analyzed by size-exclusion chromatography (Figure 2). Both nanobodies formed ternary complexes with Nb74, whose intracellular binding site had previously been established structurally (Errasti-Murugarren et al. 2019). In contrast, no ternary complex formation was detected for the extracellular pair Nb58–Nb78 or the intracellular pair Nb71–Nb74. These results support the assignment of Nb58 and Nb78 to the opposite face of the transporter relative to Nb74 and demonstrate that nanobodies recognizing opposite faces of BasC can bind simultaneously. The incompatibility observed for the same-side pairs is consistent with overlapping epitopes and/or steric interference.

**Figure 2.**
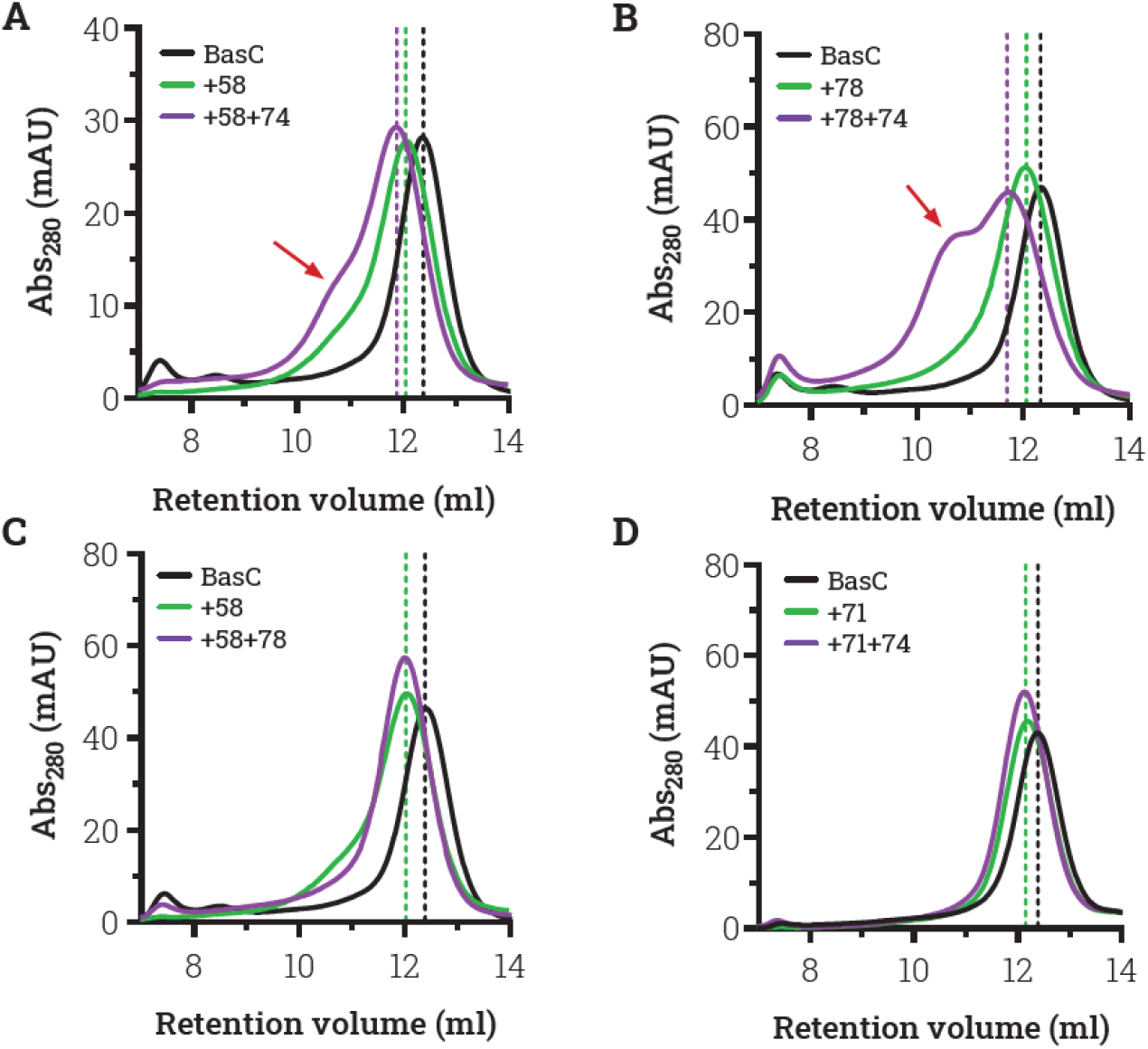
Assessment of nanobody compatibility by size-exclusion chromatography. Representative SEC profiles of DDM-solubilized BasC alone (black), BasC incubated with a first nanobody (green), and the complex following addition of a second nanobody (purple), analyzed on a Superdex 200 10/300 GL column. Each nanobody was added at a BasC molar ratio of 1:1.5. **(A, B)** Compatible pairs Nb58– Nb74 (A) and Nb78–Nb74 (B) show a shift towards lower retention volumes following addition of the second nanobody, consistent with ternary-complex formation. Red arrows indicate additional earlier-eluting species. **(C, D)** Nb58–Nb78 (C) and Nb71–Nb74 (D) show no further shift following addition of the second nanobody, consistent with incompatible binding under the conditions tested. Colour-matched vertical dashed lines indicate the mean retention volumes of the corresponding species obtained from experiments with either order of nanobody addition. Profiles are shown over the relevant retention-volume range.

### Structural basis for simultaneous nanobody binding from opposite sides of BasC

To facilitate structural localization of extracellular nanobodies, BasC was co-crystallized with Nb53 and Nb58. Addition of Nb58 improved crystal packing and enabled determination of the overall architecture of the ternary complex (Supplementary Figure S6).

The crystals diffracted to 6.1 Å resolution. Molecular replacement using the previously reported BasC structure (Errasti-Murugarren et al. 2019) revealed two nanobodies bound simultaneously on opposite sides of the transporter. However, the limited resolution did not allow atomic modelling of the binding interfaces or unambiguous identification of each nanobody from the electron density alone.

The published cryo-EM structure of the BasC–Nb53 complex (Fort et al. 2025) fitted well into the crystal density (Supplementary Figure S6E), strongly suggesting that the nanobody bound to the intracellular region surrounding intracellular loop 4, between TM8 and TM9, corresponds to Nb53. Accordingly, the nanobody bound on the opposite side of the transporter was assigned as Nb58, which is consistent with the transport-sidedness experiments identifying Nb53 and Nb58 as intracellular and extracellular binders, respectively (Figure 3). The density also indicates that BasC adopts the same inward-open conformation observed in the BasC–Nb53 cryo-EM structure (Supplementary Figure S6E), suggesting that Nb58 recognizes this conformation without inducing a major conformational change in the transporter.

**Figure 3.**
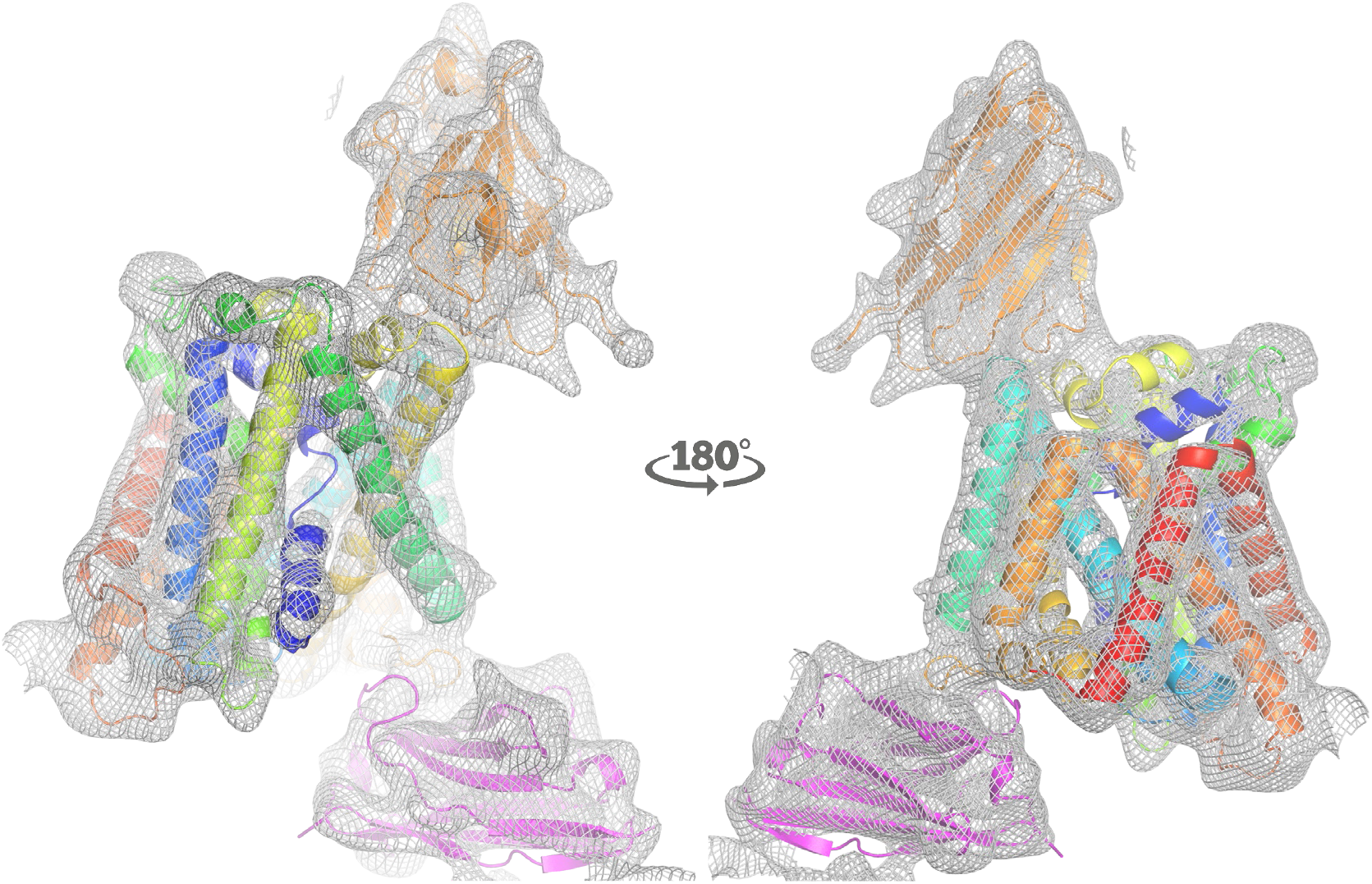
Overall architecture of the BasC–Nb53–Nb58 complex at 6.1 Å resolution. Two views related by a 180° rotation show the crystallographic electron-density map as a grey mesh, with the fitted models shown as cartoons. BasC is colored by sequence position, Nb53 in pink and Nb58 in orange. The map supports simultaneous nanobody binding to opposite faces of the transporter, with Nb53 assigned to the intracellular face and Nb58 to the extracellular face.

### AlphaFold3-based structural classification of BasC nanobody epitopes

To assess the structural distribution of the BasC nanobody epitopes, the 29 BasC–nanobody complexes were modelled with AlphaFold3 using sequence-only inputs and with templates disabled. Despite the absence of membrane lipids or sidedness restraints, the predictions placed the nanobodies on defined regions of either the intracellular or extracellular side of BasC. BasC adopted a highly similar global conformation across all 29 AF3 models. Superposition against the BasC chain of the Nb74 prediction yielded Cα RMSD values of 0.17– 0.67 Å for the remaining 28 models. The high-confidence models, defined by ipTM >0.75, were therefore mapped onto BasC and colored according to their predicted epitope group (Figure 4). This analysis revealed several recurrent intracellular epitopes. The nanobodies with the most internal binding mode formed a Nb71-like group, located deep within the intracellular vestibule (Figure 4A-C). A second group, including Nb53, Nb55 and Nb60 predictions, bound a related inward-facing region (Figure 4D), whereas Nb62 and Nb66 contacted a more lateral intracellular epitope involving TM12, TM3 and the TM2–TM3 loop (Figure 4F). A further group, related to the crystallographic Nb74 epitope, mapped to a more external intracellular region. In contrast, most extracellular nanobodies were associated with low-confidence AF3 predictions, with Nb58-like binding being poorly recovered. Nb78 was the only high-confidence extracellular prediction, although its binding mode appeared atypical and not primarily CDR3-driven (Figure 4 E).

**Figure 4.**
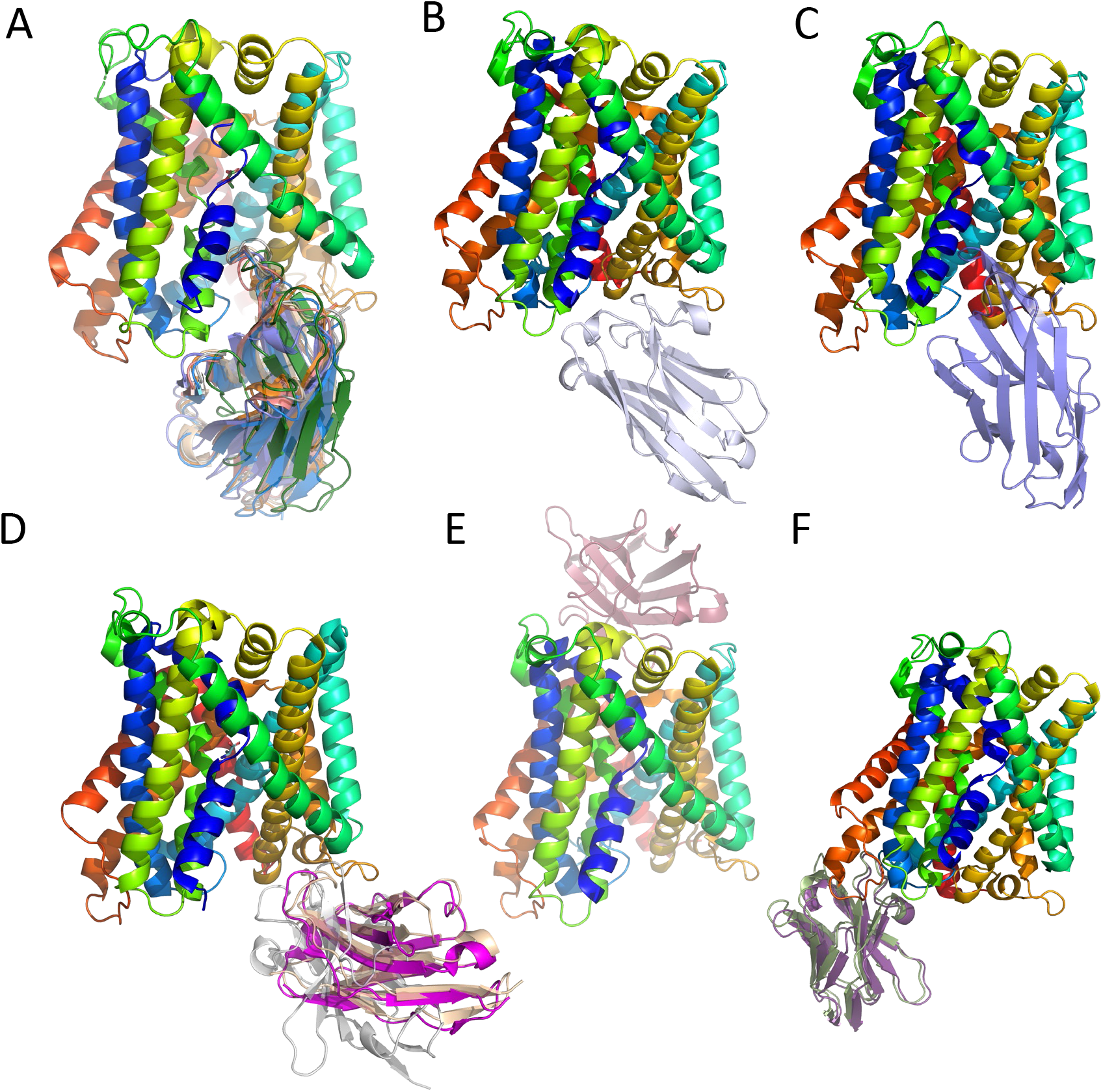
Predicted nanobody epitope distribution from high-confidence AlphaFold3 models (ipTM > 0.75). Models were grouped according to their predicted binding location and orientation and superposed using the BasC chains. **(A)** Nb71-like cluster comprising Nb56, Nb57, Nb63, Nb64, Nb69, Nb70 and Nb71. **(B, C)** Nb67 and Nb73 bind within the intracellular cavity but adopt distinct poses. **(D)** Nb53, Nb55 and Nb60 recognize a similar intracellular region: Nb60 closely reproduces the Nb53 binding mode, whereas Nb55 adopts a slightly different pose. **(E)** Nb78, the only extracellular binder among the high-confidence models. **(F)** Nb62 and Nb66 share a distinct predicted intracellular epitope. BasC is colored by sequence position, with the extracellular face oriented upwards. Superposed nanobodies represent separate complexes and do not imply simultaneous binding.

Comparison with the available experimental BasC–nanobody structures supported the interpretation of the high-confidence intracellular models (Supplementary Figure S7). AF3 reproduced the binding modes of the intracellular nanobodies Nb53, Nb71, and Nb74 (Fort et al. 2025; Errasti-Murugarren et al. 2019), with Cα RMSD values of 0.51, 1.90, and 0.51 Å, respectively, after superposition of the BasC chains. By contrast, AF3 failed to recover the extracellular Nb58 interaction (Supplementary Figure S7); however, the Nb58 prediction had a low ipTM (0.16), indicating that the confidence score correctly flagged this model as unreliable. Importantly, the predictions were generated without templates. Thus, although the previously published BasC–Nb74 structure could in principle have been included in the pre-2021 structural data available during AF3 training, the agreement with Nb71 cannot be attributed to template retrieval during inference. In contrast, the BasC–Nb53 structure was released in 2025 and therefore could not have contributed to AF3 training or template information under the reported 2021 cutoff.

Overall, AF3 generated high-confidence interaction predictions for 16 of the 29 nanobodies. Ten of these models mapped to the intracellular cavity, including a prominent Nb71-like cluster comprising seven nanobodies, while the remaining models identified alternative binding regions. These results reveal a marked convergence of predicted epitopes toward the intracellular face of BasC, consistent with the functional sidedness assignments.

## Discussion

The BasC nanobody repertoire showed a pronounced bias towards the intracellular face, with recurrent predicted binding modes within the inward-facing cavity. The agreement between functional sidedness measurements and experimentally supported AF3 models suggests that this cavity is a major target within the repertoire, despite the diversity of CDR3 sequences.

Combining transport inhibition with independent biochemical and structural evidence strengthened the sidedness assignments. Compatibility with the intracellular binder Nb74 supported the extracellular localization of Nb58 and Nb78. The BasC–Nb53–Nb58 crystal structure further supported this assignment for Nb58, showing the two nanobodies bound on opposite faces of the transporter.

The influence of target conformation on nanobody selection has been demonstrated for LacY, where nanobodies raised against an outward-open mutant stabilized outward-facing conformers of the wild-type transporter (Smirnova et al. 2014). Similarly, in vitro sybody selections against SLC and ABC transporters used ligands to favor defined target states (Zimmermann et al. 2018). These studies illustrate how antigen conformation and selection conditions can shape the recovered binder repertoire and how nanobodies can stabilize specific protein conformations (Uchański et al. 2020).

The strong intracellular bias of the repertoire raises questions about the structural determinants of nanobody recognition. Among the 16 high-confidence AF3 predictions, ten positioned the nanobody within the intracellular cavity, including a prominent Nb71-like cluster comprising seven nanobodies. This distribution is consistent with the predominance of intracellular inhibitors identified experimentally and suggests that the inward-facing cavity represents a particularly favorable region for nanobody binding. The cavity constitutes a large and structurally complex surface that may accommodate different complementarity-determining regions and support multiple partially overlapping epitopes. BasC was randomly oriented in the proteoliposomes used for immunization (Bartoccioni et al. 2019), indicating that the starting preparation did not selectively expose one face of the transporter. The observed enrichment may therefore reflect differences in cavity geometry, accessibility, or conformational stability. Nevertheless, conformational selection during immunization and nanobody screening may also have contributed to this distribution.

All predictions were generated with structural templates disabled. The predicted binding modes of Nb53, Nb71, and Nb74 closely matched the corresponding experimental structures, whereas the binding mode of the extracellular nanobody Nb58 was not correctly recovered and received a low confidence score. BasC adopted a similar global conformation in both high- and low-confidence predictions. However, these results do not exclude the possibility that some nanobodies recognize alternative conformational states not recovered by AF3. These observations are consistent with previous studies showing that AlphaFold-based approaches can accurately predict a subset of antibody–antigen interactions, although their performance remains dependent on the specific complex and the confidence of the predicted interface (Yin and Pierce 2024; Abramson et al. 2024). Importantly, confidence in the predicted binding mode should not be interpreted as a measure of binding strength. In our repertoire, no correlation was observed between ipTM and experimental binding affinity, indicating that structural prediction confidence and nanobody affinity provide distinct and complementary information.

Beyond the prediction of individual epitopes, AF3 enabled the identification of recurrent binding modes among nanobodies with divergent CDR3 sequences. This structural classification revealed groups of nanobodies targeting similar regions of BasC despite limited sequence similarity, as well as alternative epitopes distributed across the intracellular surface. Such information may be particularly useful for selecting nanobody combinations, since predicted epitope overlap can suggest potential competition, whereas spatially separated binding regions may identify candidates for simultaneous binding. However, structural proximity alone cannot establish compatibility, because conformational changes, steric constraints, and allosteric effects may also influence complex formation. The experimentally observed compatibility of Nb53 or Nb74 with extracellular nanobodies illustrates the value of combining structural predictions with biochemical validation.

An additional implication of our results concerns the selection of nanobodies for structural studies. Notably, Nb53, Nb71, and Nb74, which previously yielded the most informative crystallographic, cryo-EM, and smFRET data, were all included among the high-confidence AF3 predictions. This retrospective observation suggests that confidence-filtered structural prediction could help identify promising candidates early in the characterization process. Nevertheless, the failure to correctly model Nb58, despite its strong experimental inhibition and utility for stabilizing the ternary complex, illustrates an important limitation. Low-confidence predictions should not be used to exclude experimentally relevant nanobodies. Rather, AF3 may be most effective as a complementary screening tool that reduces the number of candidates requiring extensive experimental characterization while preserving structural and functional diversity.

Several limitations should be considered. Functional sidedness assignments remained undetermined for nanobodies showing no significant inhibition or ambiguous effects across the two transport assays. The low-resolution BasC–Nb53–Nb58 structure defines the overall arrangement of the complex but does not resolve the extracellular binding interface at atomic resolution. Furthermore, most AF3-predicted epitopes remain experimentally unvalidated, and high-confidence predictions do not necessarily imply strong binding or effective transport inhibition. Despite these limitations, the convergence of functional measurements, compatibility experiments, experimental structures, and structural predictions supports the overall classification of the repertoire. Our findings suggest that, for transporters with asymmetric substrate affinities, integrating transport-based sidedness determination with confidence-filtered AF3 modeling can provide an effective strategy for mapping nanobody epitopes and prioritizing structurally informative binders.

## Methods

### BasC expression and membrane preparation

BasC and BasC–GFP were expressed and purified essentially as previously described (Errasti-Murugarren et al., 2019; Bartoccioni et al., 2019). Briefly, Escherichia coli BL21 Star (DE3) cells transformed with the corresponding construct were grown in LB medium supplemented with 0.1 mg ml™1 ampicillin at 37 °C and 240 rpm. Cultures were induced at an OD600 of 0.5 with 0.1 mM isopropyl β-D-1-thiogalactopyranoside and incubated overnight at 37 °C. Cells were collected at 5,000 × g for 30 min at 4 °C and washed in high-salt buffer containing 20 mM Tris-HCl, pH 7.4, and 350 mM NaCl. Cell pellets were resuspended in the same buffer supplemented with DNase (1:500) and one cOmplete EDTA-free protease inhibitor tablet per 50 ml, and disrupted by four passes at 35 kpsi. Unbroken cells were removed by centrifugation at 20,000 × g for 1 h at 4 °C, and membranes were recovered from the supernatant by centrifugation at 200,000 × g for 2 h at 4 °C. Membranes were resuspended in 20 mM Tris-HCl, pH 7.4, 150 mM NaCl (standard Tris buffer) and 10% (v/v) glycerol, snap-frozen in liquid nitrogen and stored at −80 °C.

### BasC purification

Membranes were diluted to 3 mg ml™1 in standard Tris buffer supplemented with either 1% (w/v) n-dodecyl-β-D-maltoside (DDM) or 2% (w/v) n-decyl-β-D-maltoside (DM), and solubilized for 1 h at 4 °C. Insoluble material was removed by centrifugation at 200,000 × g for 2 h. The supernatant was incubated for at least 3 h at 4 °C with Ni^+^–NTA Superflow resin (0.5 ml resin per 10 ml solubilized membranes). All subsequent buffers contained 0.05% (w/v) DDM or 0.17% (w/v) DM, as appropriate. The resin was washed sequentially with 20 mM Tris-HCl, pH 7.4, and 150 mM NaCl supplemented with 10, 20 and 30 mM imidazole. BasC–GFP was eluted with the same buffer containing 300 mM imidazole and concentrated using a 100-kDa molecular-weight-cut-off centrifugal filter. For tag-free BasC, the resin was equilibrated in 20 mM Tris-HCl, pH 7.4, 150 mM NaCl, 1 mM dithiothreitol and 1 mM EDTA, supplemented with the corresponding detergent, and incubated overnight at 4 °C with HRV 3C protease. Cleaved BasC was collected and concentrated using a 50-kDa molecular-weight-cut-off centrifugal filter. DDM-purified protein (0.05% DDM) was used for binding and transport assays, whereas DM-purified BasC (0.17% DM) was used for structural studies unless otherwise stated.

### Nanobody generation, selection and sequence analysis

Nanobodies against BasC were generated using purified transporter reconstituted into liposomes as antigen, following established procedures for the generation of nanobodies for structural biology (Pardon et al., 2014). The selection yielded 73 ELISA-positive clones. Nanobody sequences were analysed and grouped into 27 families, primarily according to CDR3 sequence similarity, and representative clones were selected to obtain the panel of 29 nanobodies examined in this study. Nanobody sequences and family assignments are provided in Supplementary Table S1 and Supplementary Figure S2.

### Nanobody expression and purification

Nanobodies were expressed in Escherichia coli WK6 (Su−) cells. Starter cultures were grown overnight at 37 °C and 190 rpm in LB medium supplemented with 0.1 mg ml−1 ampicillin, 2% (w/v) glucose and 1 mM MgCl2. Terrific Broth cultures supplemented with 0.1 mg ml−1 ampicillin, 0.1% (w/v) glucose and 2 mM MgCl2 were inoculated with the starter cultures and grown at 37 °C and 190 rpm to an OD600 of 0.7–1.1. Expression was induced with 1 mM isopropyl β-D-1-thiogalactopyranoside and continued overnight at 28 °C.

Cells were harvested at 4,500 × g for 30 min at 4 °C and subjected to osmotic shock. Cell pellets were resuspended in TES buffer (200 mM Tris-HCl, 0.5 mM EDTA and 500 mM sucrose) (15 ml per litre of initial culture at an OD600 equivalent to 25), stirred for at least 1 h at 4 °C, diluted with 30 ml of one-quarter-strength TES buffer and stirred for a further 45 min. The periplasmic fraction was recovered by centrifugation at 15,000 × g for 30 min and incubated for at least 3 h at 4 °C with Ni2+–NTA Superflow resin. A second osmotic-shock extraction was performed overnight. The resin was washed with 10 mM imidazole wash buffer, and nanobodies were eluted with 300 mM imidazole. The buffer was exchanged and the proteins were concentrated with 3-kDa molecular-weight-cut-off centrifugal filters, snap-frozen in liquid nitrogen and stored at −80 °C.

### Surface plasmon resonance

Nanobody binding to BasC was analyzed by surface plasmon resonance using a Biacore T100 instrument. To avoid detergent during ligand immobilization, His-tagged nanobodies (3–20 μg ml−1) were captured on a CM5 sensor chip coated with an anti-His antibody using a His Capture Kit. Nanobodies were immobilized to approximately 200 resonance units by injection for 20 s at 5 μl min−1. DDM-purified BasC was used as analyte at concentrations between 2.5 and 500 nM in running buffer containing phosphate-buffered saline and 0.05% (w/v) DDM. The surface was regenerated with 10 mM HCl in running buffer. Measurements were performed in the absence or presence of 4 mM L-alanine as indicated. Association and dissociation rate constants, equilibrium dissociation constants and maximum response values were obtained from the sensorgrams. The conditions used for the systematic comparison in Table 1 are specified in its legend.

### Size-exclusion chromatography and nanobody compatibility assays

BasC purified in DM or DDM was incubated for 1 h at 4 °C with the selected nanobody at a 1:1.5 BasC:nanobody molar ratio. For compatibility experiments, the second nanobody was then added at the same ratio and the mixture was incubated for an additional hour at 4 °C. A sample containing the same amount of BasC without nanobody was analysed in parallel. Samples were passed through 0.22-μm centrifugal filters and loaded onto a Superdex 200 10/300 GL column connected to an ÄKTA system. Elution was monitored at 280 nm and 250-μl fractions were collected for SDS–PAGE analysis when required. Simultaneous shifts in the BasC elution profile and the presence of both nanobodies in the complex-containing fractions were used to assess ternary-complex formation.

### Reconstitution of BasC into proteoliposomes

BasC was reconstituted into proteoliposomes essentially as previously described (Bartoccioni et al., 2019). Escherichia coli total polar lipid extract was dissolved in chloroform at 50 mg ml−1. Lipids were dried under a stream of nitrogen to form a thin film and maintained under nitrogen for 3 h at room temperature. The lipid film was resuspended by sonication in standard Tris buffer to 40 mg ml−1. Purified BasC–GFP was mixed with the lipids at a 1:100 protein:lipid mass ratio. Octyl glucoside was added to 1.25% (w/v), and the mixture was incubated for 30 min at 4 °C. Detergent was removed by dialysis for 40 h at 4 °C against 100 volumes of standard Tris buffer. Proteoliposomes were recovered, snap-frozen in liquid nitrogen and stored at −80 °C.

### Radiolabelled substrate influx and efflux assays

Influx and efflux were measured in BasC–GFP proteoliposomes using radiolabelled L-serine. BasC inserts into proteoliposomes with random orientation and displays different apparent substrate affinities at its extracellular and intracellular faces. At the low substrate concentrations used here, influx and efflux therefore preferentially report transport by right-side-out and inside-out BasC molecules, respectively.

For influx measurements, proteoliposomes were loaded with 4 mM L-alanine by six freeze– thaw cycles. Control liposomes were subjected to the same procedure without amino acid. After incubation on ice for 30 min, proteoliposomes were collected at 100,000 × g for 1 h at 4 °C and resuspended in standard Tris buffer to one-third or one-half of the initial volume, depending on the preparation. Transport was initiated by mixing 20 μl of proteoliposomes with 180 μl of standard Tris buffer containing 10 μM L-serine and 0.5 μCi L-[3H]-serine. After 5 s, transport was stopped by adding 3 ml of ice-cold standard Tris buffer supplemented with 10 mM L-serine.

For efflux measurements, proteoliposomes were loaded with 10 μM L-serine and 0.5 μCi L-[3H]-serine per assay point by six freeze–thaw cycles. After incubation on ice and ultracentrifugation as described above, efflux was initiated by mixing 20 μl of loaded proteoliposomes with 180 μl of standard Tris buffer containing 4 mM L-alanine. After 30 s, transport was stopped by adding 3 ml of ice-cold standard Tris buffer.

Samples were filtered through 0.45-μm membrane filters and washed twice with 3 ml of the corresponding ice-cold stop buffer. Filters were dried at room temperature, immersed in scintillation fluid and counted for 3 min. For influx, time-zero controls were generated by adding stop solution before the proteoliposomes, and transport measured in unloaded proteoliposomes was subtracted from that measured in L-alanine-loaded proteoliposomes. Influx values were normalized to the amount of reconstituted BasC–GFP determined by SDS– PAGE and expressed as pmol radiolabelled amino acid μg−1 BasC s−1. For efflux, loss of intraliposomal radioactivity was calculated relative to time-zero controls. Additional 30-s controls without external L-alanine were included throughout the experiment. Efflux values were expressed relative to the corresponding untreated sample.

### Nanobody-mediated transport inhibition and sidedness assignment

Nanobodies were added exclusively to the extraliposomal solution and preincubated with the proteoliposomes for 20 min before initiating transport. The nanobody concentration was 15 μM for the influx screen and 5 μM for the efflux screen. All nanobodies were measured in duplicate; a third measurement was obtained for nanobodies showing inhibition in either of the first two measurements. Transport activity was expressed relative to proteoliposomes incubated without nanobody.

### Crystallization of the BasC–Nb53–Nb58 complex

For crystallographic studies, freshly purified BasC in DM was concentrated to 6 mg ml−1 using a 100-kDa molecular-weight-cut-off centrifugal filter. Nanobodies were added at a 1:1.2 BasC:nanobody molar ratio, and complexes were incubated for 1 h at 4 °C and purified on a Superdex 200 10/300 GL column equilibrated in DM buffer at 0.5 ml min−1. Complex-containing fractions were pooled and concentrated to approximately 2 mg ml−1. The presence of BasC and the nanobodies was verified by SDS–PAGE.

BasC–Nb53–Nb58 was screened in membrane-protein crystallization conditions using sitting- or hanging-drop vapor diffusion at 4 and 20 °C. Protein and reservoir solutions were mixed at a 1:1 volume ratio. Crystals were obtained in 50 mM ADA, pH 6.8, 6.4% (w/v) PEG MME 2000 and 17.1% (w/v) PEG 3000 and were optimized around this condition. Crystals were cryoprotected in reservoir solution and flash-cooled in liquid nitrogen.

### X-ray data collection and structural analysis

Diffraction data for the BasC–Nb53–Nb58 complex were collected at the ID23-1 beamline of the European Synchrotron Radiation Facility (Grenoble, France). Crystals were aligned using tilted loops with automatic re-centring to reduce the effect of anisotropy. The complete dataset was processed with autoPROC to a resolution of 6.14 Å in space group H3, with unit-cell dimensions a = b = 357.5 Å and c = 74.1 Å, and α = β = 90° and γ = 120°. The structure was solved by molecular replacement using the published BasC–Nb74 structure (Errasti-Murugarren et al., 2019) and a nanobody model lacking CDRs and side chains. Two BasC molecules and two copies of each nanobody were present in the unit cell. Nanobody assignment was guided by the transport-sidedness results and by comparison with the published BasC–Nb53 cryo-EM structure (Fort et al., 2025). Owing to the limited resolution, the structure was used to define the overall architecture and relative location of the nanobodies rather than to derive atomic details of the binding interfaces.

### AlphaFold3 modelling of BasC–nanobody complexes

Structural models of the BasC–nanobody complexes were generated with AF3 to explore the possible epitope distribution of the complete nanobody panel. The amino acid sequence of BasC and the sequences of each individual nanobody were used as input, generating one independent prediction for each BasC–nanobody pair. A total of 29 complexes were modelled. Only protein sequences were provided as input, and structural templates were explicitly disabled by setting the template field to an empty list for each protein chain. No membrane lipids, membrane restraints, epitope restraints or sidedness information were included in the predictions.

For each BasC–nanobody complex, five random seeds were used, with five diffusion samples per seed, resulting in 25 predicted models per complex. The final model selected for analysis was the top-ranked AF3 prediction according to the model ranking score. Prediction confidence was evaluated using the AF3 summary confidence metrics, with particular emphasis on the interface predicted template modelling score (ipTM) and the inter-chain predicted aligned error (PAE). Models with high ipTM values and low inter-chain PAE were considered high-confidence interface predictions, whereas models with low ipTM values were interpreted cautiously and were not used for definitive epitope assignment.

For structural comparison, the BasC chains from the predicted models were superposed, and the relative positions of the nanobodies were inspected to identify recurrent epitope groups. High-confidence models were classified according to their predicted binding region on BasC, including intracellular vestibule-binding nanobodies, lateral intracellular binders, Nb74-like binders and extracellular binders. The AF3 predictions were further compared with the experimentally determined BasC–nanobody structures available for Nb53, Nb71, Nb74 and Nb58 to assess whether the predicted binding modes were consistent with the experimentally observed epitopes.

## Supporting information

Suppementary tables and figures

## Data availability

The nanobody sequences are provided in the supplementary material and will also be deposited in the NanoSaurus knowledge base (https://www.nanosaurus.org/).

Raw X-ray diffraction data collected at ESRF beamline ID23-1 under proposal MX-2356 (BAG BARCELONA) are available as part of the session dataset at doi: 10.15151/ESRF-ES-771376408.

Processed crystallographic data for the BasC–Nb53–Nb58 complex at 6.1 Å resolution are available in Mendeley Data at doi: 10.17632/4hyrf9x778.1

The top-ranked AF3 models of the 29 BasC–nanobody complexes are available in Mendeley Fort Baixeras, Joana (2026), “Nanobody-BasC Complexes AlphaFold3 predictions”, Mendeley Data, V1, doi: 10.17632/sbf38w6bhr.1

## Acknowledgments and funding

This work was supported by the Spanish national research grants PID2021-122802OB-I00 and PID2024-157999OB-I00 (PI M.P.). The authors thank INSTRUCT-ERIC (PID1176), part of the European Strategy Forum on Research Infrastructures (ESFRI), and the Research Foundation – Flanders (FWO) for their support to the Nanobody discovery. The authors also thank Nele Buys for technical assistance during Nanobody discovery. The authors thank Joan Pous from the joint IBMB-CSIC/IRB Barcelona Automated Crystallography Platform (PAC) for assistance with crystallization experiments. They acknowledge also the European Synchrotron Radiation Facility (ESRF), Grenoble, France, for the provision of synchrotron radiation facilities under proposal MX-2356, and thank Montserrat Soler-López and the staff of beamline ID23-1 for assistance with X-ray diffraction data collection. A.N.-A. was supported by an FI-AGAUR fellowship. IRB Barcelona is a member of the CERCA Programme of the Generalitat de Catalunya.

## Notes

https://doi.esrf.fr/10.15151/ESRF-ES-771376408

https://doi.org/10.17632/4hyrf9x778.1

https://doi.org/10.17632/sbf38w6bhr.1

### Competing Interest Statement

The authors have declared no competing interest.

