## Supplementary material for "Experimental and computational mapping reveals preferential nanobody targeting of the intracellular cavity of the amino acid transporter BasC": Suppementary tables and figures

**Supplementary Table S1 | Amino acid sequences of the BasC nanobody repertoire.** Full-length VHH amino acid sequences of the 29 nanobodies selected against BasC are listed together with their corresponding CDR3 sequences and original clone reference identifiers. C-terminal purification and affinity His-tag are not included. CDR3 regions were defined from the conserved cysteine preceding the variable loop to the conserved framework tryptophan immediately downstream of the CDR3.

| Name | Original reference ID | VHH amino-acid sequence | CDR3 amino-acid sequence | CDR3 length (aa) |
|---|---|---|---|---|
| Nb50 | CA11050 | QVQLVESGGGLVQPGGSLRLSCAASGRAFSDYRMGWFRQAPGKEREFVAAITWSWSTTYYADSVKGRFTISRDNAKNTVYLQMSSLKPEDTAVYFCAAQYLADYYTDGGEYKYWGQGTQVTVSS | AAQYLADYYTDGGEYKY | 17 |
| Nb51 | CA11051 | QVQLVESGGGLVQPGNSLRLSCAASGRALHDYRMGWFRQAPGKEREFVAAIGWSSGTPYYADSVKGRFAISRDSSKNTVFLQMNSLKPEDTAVYSCAAQYWGDYYSDGAEYKYWGQGTQVTVSS | AAQYWGDYYSDGAEYKY | 17 |
| Nb52 | CA11052 | QVQLVESGRGVVQAGGSLRLSCAASGRTFSSYAMGWFRQAPGKEREFVAFISGSGISTRYADSVQGRFTITRDNAKNAVYLQMNSLLPEDTAVYYCAADQKHFQTRASNYHYWGQGTQVTVSS | AADQKHFQTRASNYHY | 16 |
| Nb53 | CA11053 | QVQLVESGGGLVQAGGSLRLSCAASGIAFSRMSMAWYRQDPGKQRALVARITNDGSTYYDDSVKGRFTISRDNAKNTVHLQMNSLKPEDTAVYYCNAQLVAWSENYWGQGTQVTVSS | NAQLVAWSENY | 11 |
| Nb54 | CA11054 | QVQLVESGGGLVQAGGSLTLSCAASGRSFSSYRMGWFRQPPGKEREFVATISRSGGGTYYADSVKGRFTISRDNAKNMVYLLMNTLKPEDTAVYYCAADYTLYSDTYYSSDSYDYWGQGTQVTVSS | AADYTLYSDTYYSSDSYDY | 19 |
| Nb55 | CA11055 | QVQLVESGGGLVQPGDSLRLSCAAPGRTLSGRYMGWFRQAPGKEREFVAAFNWGSIEYADSVKGRFIISRDNAKNTVYLQMNSLKPEDTAVYYCAARWAYSGTYDAVAGYDYWGQGTQVTVSS | AARWAYSGTYDAVAGYDY | 18 |
| Nb56 | CA11056 | QVQLVESGGGLVQPGGSLRLSCAASGSSTSGINAMAWYRQAPGKQREFVAIVTSGGSTSYADSVKGRFTISADNAKNTVYLQMNSLKPEDTAVYYCYARSRRLVTTYDYWGQGTQVTVSS | YARSRRLVTTYDY | 13 |
| Nb57 | CA11057 | QVQLVESGGGLVQAGGSLRLSCAASGSFFSITAMGWYRQAPGKQRELVATITSGGRTDYADSMKGRFTISGDNAKNTVYLQMNSLKPEDTAVYYCKAQGRMSWTLTYNDFWGQGTQVTVSS | KAQGRMSWTLTYNDF | 15 |
| Nb58 | CA11058 | QVQLVESGGGLVPAGGSLRLSCAASGRTFSIYAMAWFRQPPGKEREFVARIDWSGGRIAYADSVEGRFTISRDNSAKNTWYLQMNSLKPDDTAVYYCASARTGWSTSQDSYDYWGEGTQVTVSS | ASARTGWSTSQDSYDY | 16 |
| Nb59 | CA11059 | QVQLVESGGGLVQAGGSLTLSCTASGSIFSNSAMGWYRQAPGKQREFVARIASVGSTNYADSVKGRFTISRGNAKNTVYLQMSSLKPEDTAVYFCNAAQRVSPITLHTVDYWGQGTQVTVSS | NAAQRVSPITLHTVDY | 16 |
| Nb60 | CA11060 | QVQLVESGGGLVQAGGSLRLYCAGSERTFDDYTMGWFRQAPGKPRELVAAITRASDIYYRDSVKGRFSIARENAKNTLYLQMNSLKPEDTATYYCAASESGTGWDHSGVRWGHGTQVTVSS | AASESGTGWDHSGVR | 15 |
| Nb61 | CA11061 | QVQLVESGGGLVQAGGSLRLSCAASGRTRGSYAMGWFRQAPGKERDFVAAISSSGSSTYYAHSVKGRFTISRDNAKSTVYLQMNSLKPEDTAVYYCAADYGLFSDLSDAKEYDYWGQGTQVTVSS | AADYGLFSDLSDAKEYDY | 18 |
| Nb62 | CA11062 | QVQLVESGGGLVQAGGSLRLSCAASGRTFSTYNMGWFRQAPGKEREFVAAIRKSGSSTYYTSSVKGRFTITRDNAANTVYLQMNSLKPEDTAVYYCAADFALFSDLRDSREYDYWGQGTQVTVSS | AADFALFSDLRDSREYDY | 18 |
| Nb63 | CA11063 | QVQLVESGGGLVQPGGSLRLSCAASGNIDVITGMGWYRQAPGKQRELVALITRGGRINYAASVRGRFTISSDNAKNTVYLQMNSLEPEDTAVYYCQLTQRGRFSLSSTSDYWGQGTQVTVSS | QLTQRGRFSLSSTSDY | 16 |
| Nb64 | CA11064 | QVQLVESGGGLVQPGGSLRLSCAVSGSTFVINVMSWYRQAPGNQRELVASITSSGGTNYADSVKGRFTISRDNAENYLYLQMNSLKPEDTAVYYCNAEGRYRGSLIRPFPIWGQGTQVTVSS | NAEGRYRGSLIRPFPI | 16 |
| Nb65 | CA11065 | QVQLVESGGGYVQAGDSLRLSCAASGRTFSTYFMGWFRQAPGKEREFVAAISESRSSTFYADSVKGRFTISRDNTKSTGYLQMASLKTEDTAVYFCAAGRGPFKMLTMRAAYDYWGQGTQVTVSS | AAGRGPFKMLTMRAAYDY | 18 |
| Nb66 | CA11066 | QVQLVESGGGLVQAGGSLRLSCAASGRTFSSYTMGWFRQAPGKEREFVAAIKSSDRSTSYAYSVKGRFTISRDNAKNTVYLQMDSLKPEDTAVYYCAATSRLFPVRSDENEYDYWGQGTQVTVSS | AATSRLFPVRSDENEYDY | 18 |
| Nb67 | CA11067 | QVQLVESGGGGVQAGDSLRLSCAASGRTFSSYGMGWFRQAPGKEREFVAGINWNGGSTSYADSVKGRFTISRDNAKNTVYLQMNNLKPADTAVYYCARSREMWYSSALHDHDYWGQGTQVTVSS | ARSREMWYSSALHDHDY | 17 |
| Nb68 | CA11068 | QVQLVESGGGLVQAGGSLRLSCAASGRIFGNYVMGWFRQAPGKEREFVAAVSESGGRTYYADSVKGRFTISRDNAKNTVYLQMNSLNLEDTAVYYCAADAVRLTVRTSYNYWGQGTQVTVSS | AADAVRLTVRTSYNY | 15 |
| Nb69 | CA11069 | QVQLVESGGGSVQPGGSLRLSCAASGNIFIITGMGWYRQAPGKQRELVAQITSGGSTVYADSVKDRFTISENNAKNTVYLQMNSLQPEDTAVYYCQLSQRHYFDMGYSSDYWGQGTQVTVSS | QLSQRHYFDMGYSSDY | 16 |
| Nb70 | CA11070 | QVQLVESGGGLVQAGGSLRLSCAASGSFANINTMGWYRQAPGNQRELVAIITAGGRQNYADSVKGRFTIFGDNPKNTVYLQMNNLKPEDTAVYYCNALVRTGINWPVDYWGQGTQVTVSS | NALVRTGINWPVDY | 14 |
| Nb71 | CA11071 | QVQLVESGGGLVQAGGSLRLSCAASSDIFTITAMGWYRQAPGKQRELVASITSGGRKNYADSVKGRFTISSDNAKKTVNLQMNSLEPEDTAVYYCHVNYRPLISLDIFDAWGQGTQVTVSS | HVNYRPLISLDIFDA | 15 |
| Nb72 | CA11072 | QVQLVESGGGLVQPGGSLRLSCAASGGIFTIRSMGWYRQAPGKERELVAMITSGGLINYDDSVKGRFTISRDSAKVTVYLQMNSLRPEDTAVYYCNAELRRSIVMNYAEYWGQGTQVTVSS | NAELRRSIVMNYAEY | 15 |
| Nb73 | CA11073 | QVQLVESGGGWVQAGGSLRLSCAASGSMFGFSAMGWYRQAPGKQRELVATISSGGTTNHADSAKGRFTLTSDNNENTVYLQMNDLKPEDTAVYYCKAVARLGLGITSDYWGQGTQVTVSS | KAVARLGLGITSDY | 14 |
| Nb74 | CA11074 | QVQLVESGGGVVQAGGSLRLSCAASGRTFSSRAMGWFRQAPGEGREFVATISWSGSYTEYADSVKGRVTISRDNAKNTVYLQMNSLKPGDTAVYHCAAKNGGAASNYPNDYVYWGQGTQVTVSS | AAKNGGAASNYPNDYVY | 17 |
| Nb75 | CA11075 | QVQLVESGGRLVQAGGSLRLSCAASGRAFDSYTMAWFREAPGKEREIVGSITKRGGITNYTESVKGRFTISKDDARNTVYLQMNNLKPEDTANYICAADRRSGLIMDTVGIDYWGLGTQVTVSS | AADRRSGLIMDTVGIDY | 17 |
| Nb76 | CA11076 | QVQLVESGGGLVQAGGSLRLSCVVSGRTFGDYTYGWVRQSPGKQREFVAAINWSGTTTDYRDSVKGRLRISRDNARNTVYLQMNNLKPEDTAVYTCATPQWTKYFSSKDHFASFDYWGRGTQVTVSS | ATPQWTKYFSSKDHFASFDY | 20 |
| Nb77 | CA11077 | QVQLVESGGGLVQAGGSLRLSCTASNNMFRLKVVGWHRQAPGKQRELVASITTGGTTNVADSVKGRFTISRDNAQNMVYLQMNGLKPEDTAVYYCNADLFELGTARFPNYWGQGTQVTVSS | NADLFELGTARFPNY | 15 |
| Nb78 | CA11078 | QVQLVESGGGLVQPGGSLRLSCLASGFTLRNYPMSWVRQAPGKGLEWVAGISTYGGFTNYAESVKGRFTISRENDNNMLYLQMNSLKPEDTAVYYCAQGIDQRTVVGSSRGQGTQVTVSS | AQGIDQRTVVGSSR | 14 |

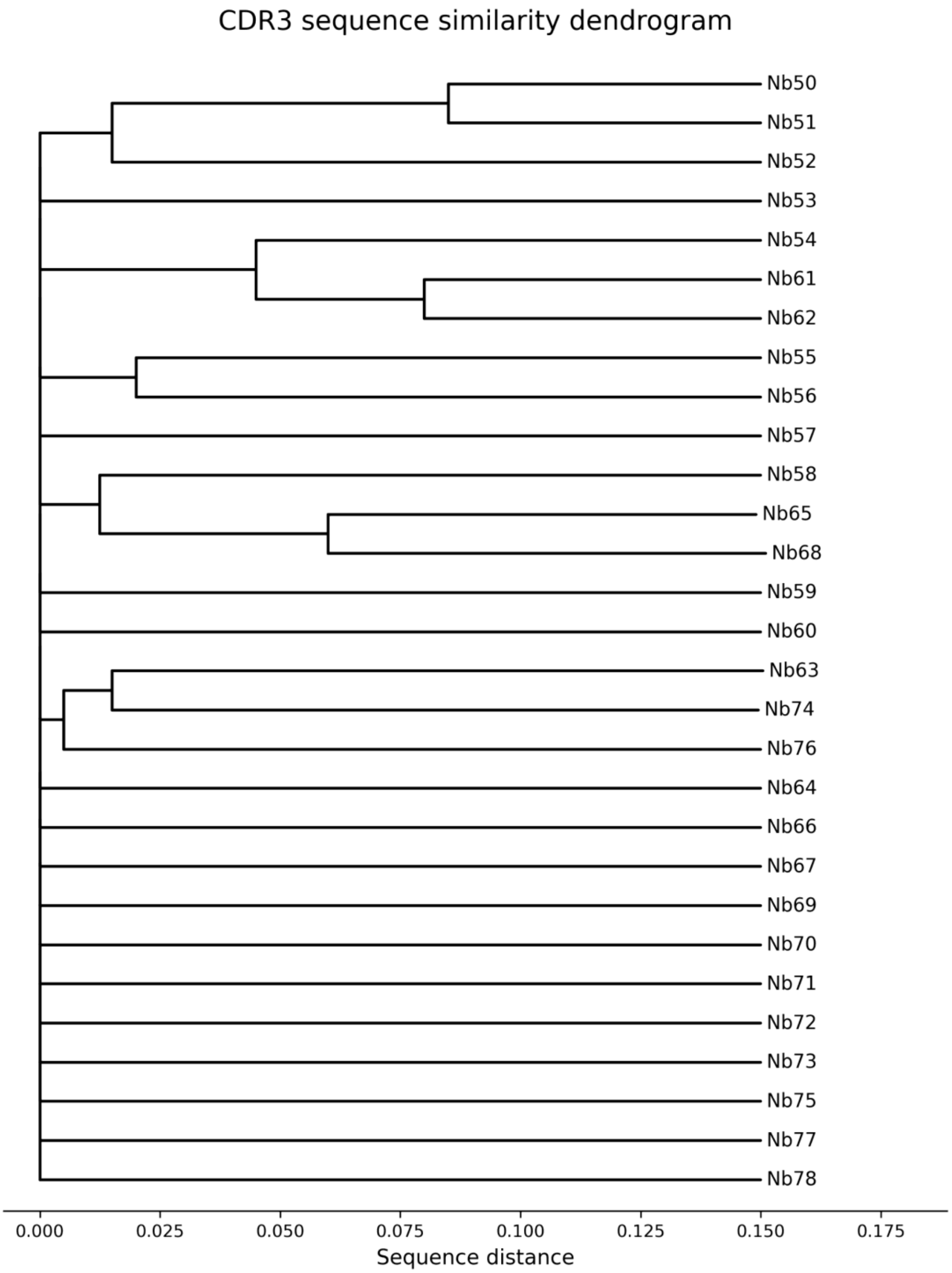


**Supplementary Figure S2 | CDR3 sequence similarity within the BasC nanobody repertoire.** Dendrogram based on the amino acid sequences of the CDR3 regions from the 29 BasC nanobodies. CDR3 sequences were aligned using T-Coffee, and sequence distances were used to generate the dendrogram. Nanobodies are indicated as Nb50–Nb78. The analysis reveals a highly diverse CDR3 repertoire, with only a limited number of closely related sequence clusters.

| Nanobody | $K_D$ apo (nM) | $R_{max}$ apo (RU) | $k_a$ apo $M^{-1} s^{-1}$ | $k_d$ apo $s^{-1}$ | $K_D$ + L-Ala (nM) | $R_{max}$ + L-Ala (RU) | $k_a$ + L-Ala $M^{-1} s^{-1}$ | $k_d$ + L-Ala $s^{-1}$ | SEC | Influx (%) | Efflux (%) |
|---|---|---|---|---|---|---|---|---|---|---|---|
| Nb50 | 4.91 | 379.0 | 4.71e+05 | 2.31e-03 | 23.10 | 928.0 | 1.10e+05 | 2.55e-03 | Yes | 69.2 | 26.2 |
| Nb51 | 36.13 | 200.0 | 6.19e+05 | 2.24e-02 | 111.00 | 518.0 | 2.64e+05 | 2.95e-02 | Yes | 86.7 | 13.9 |
| Nb52 | 34.29 | 17.0 | 1.21e+06 | 4.17e-02 | ND | 1.2 | ND | ND | ND | 78.1 | 32.7 |
| Nb53 | 26.40 | 497.0 | 2.31e+05 | 6.10e-03 | 57.80 | 762.0 | 1.26e+05 | 7.28e-03 | Yes | 96.2 | -4.5 |
| Nb54 | 72.00 | 51.0 | 7.87e+04 | 5.73e-03 | 67.80 | 53.0 | 6.58e+04 | 4.46e-03 | Yes | 88.5 | 17.4 |
| Nb55 | 101.05 | 70.0 | 1.79e+07 | 1.81e+00 | 7.12 | 10.1 | 3.34e+05 | 2.38e-03 | Yes | 75.7 | 3.0 |
| Nb56 | 26.30 | 200.0 | 1.28e+05 | 2.26e-03 | 14.00 | 835.0 | 1.71e+05 | 2.39e-03 | Yes | 95.8 | 33.0 |
| Nb57 | 216.58 | 80.0 | 2.30e+05 | 4.99e-02 | 238.00 | 71.0 | 2.03e+05 | 4.84e-02 | Yes | 85.7 | 6.1 |
| Nb58 | 28.10 | 289.0 | 9.94e+04 | 2.79e-03 | 43.10 | 717.0 | 1.67e+05 | 7.20e-03 | Yes | 13.0 | 85.0 |
| Nb59 | 248.00 | 80.0 | 6.06e+05 | 6.06e-01 | 157.00 | 36.0 | 2.26e+05 | 3.55e-02 | No | 67.0 | 15.0 |
| Nb60 | 18.60 | 260.0 | 1.62e+05 | 3.01e-03 | 86.30 | 2290.0 | 3.99e+04 | 3.45e-03 | Yes | 96.7 | 26.7 |
| Nb61 | 33.00 | 15.0 | 2.60e+05 | 8.59e-03 | ND | 5.0 | ND | ND | No | 100.4 | 71.4 |
| Nb62 | ND | 20.0 | ND | ND | ND | 2.6 | ND | ND | ND | 114.9 | 56.4 |
| Nb63 | ND | 15.0 | ND | ND | ND | 1.1 | ND | ND | No | 73.2 | 21.4 |
| Nb64 | ND | 4.0 | ND | ND | ND | 0.0 | ND | ND | No | 52.8 | -8.0 |
| Nb65 | 11.50 | 241.0 | 1.73e+05 | 2.00e-03 | 26.50 | 621.0 | 7.81e+04 | 2.07e-03 | ND | 83.1 | 29.0 |
| Nb66 | 115.00 | 50.0 | 1.00e+12 | 1.00e+05 | 16.70 | 9.1 | 2.58e+05 | 4.31e-03 | ND | 75.7 | 57.6 |
| Nb67 | 97.00 | 220.0 | 4.76e+05 | 4.65e-02 | 89.30 | 194.0 | 9.37e+05 | 8.37e-02 | ND | 78.9 | 21.7 |
| Nb68 | ND | 50.0 | ND | ND | 32.30 | 28.3 | 1.14e+05 | 3.69e-03 | ND | 113.4 | 92.1 |
| Nb69 | 24.90 | 218.0 | 1.26e+05 | 3.13e-03 | 69.30 | 332.0 | 4.14e+04 | 2.87e-03 | ND | 95.4 | 20.4 |
| Nb70 | 57.00 | 120.0 | 1.51e+05 | 7.95e-03 | 224.00 | 191.9 | 3.35e+04 | 7.51e-03 | Yes | 97.0 | 3.5 |
| Nb71 | 135.00 | 1550.0 | 2.55e+04 | 3.43e-03 | 173.00 | 1629.5 | 2.36e+04 | 4.09e-03 | Yes | 120.0 | 46.0 |
| Nb72 | 84.90 | 80.0 | 7.50e+05 | 6.30e-02 | 102.00 | 31.8 | 1.32e+05 | 1.35e-02 | ND | 76.7 | 22.9 |
| Nb73 | 119.00 | 100.0 | 2.80e+06 | 3.30e-01 | 70.40 | 42.4 | 3.41e+07 | 2.40e+00 | ND | 85.2 | 18.8 |
| Nb74 | 6.13 | 746.0 | 2.54e+05 | 1.56e-03 | 21.00 | 845.8 | 5.48e+04 | 1.15e-03 | Yes | 94.9 | 8.2 |
| Nb75 | ND | 8.0 | ND | ND | ND | 2.6 | ND | ND | ND | 99.2 | 87.8 |
| Nb76 | ND | 6.0 | ND | ND | ND | 0.0 | ND | ND | ND | 96.6 | 93.3 |
| Nb77 | 115.00 | 140.0 | 3.90e+05 | 4.48e-02 | 92.50 | 84.5 | 4.19e+08 | 3.88e+01 | Yes | 101.0 | 9.4 |
| Nb78 | 55.26 | 200.0 | 2.17e+05 | 1.20e-02 | 178.00 | 229.1 | 3.01e+04 | 5.36e-03 | Yes | 8.4 | 79.5 |

**Supplementary Table S3. Complete biochemical and functional characterization of the BasC nanobody repertoire.** Surface plasmon resonance measurements were performed using detergent-solubilized BasC in the absence of substrate (apo) and in the presence of L-alanine. The table reports the apparent equilibrium dissociation constant ($K_D$), maximum SPR response ($R_{\mathrm{max}}$), association rate constant ($k_a$) and dissociation rate constant ($k_d$) for each condition. SEC indicates whether formation of a stable BasC–nanobody complex was detected by size-exclusion chromatography. INF and EFF correspond to residual influx and efflux transport activities, respectively, expressed as percentages relative to the corresponding control condition; lower values therefore indicate stronger inhibition. All nanobodies are included, including those for which binding or kinetic parameters could not be reliably determined. ND, not determined.

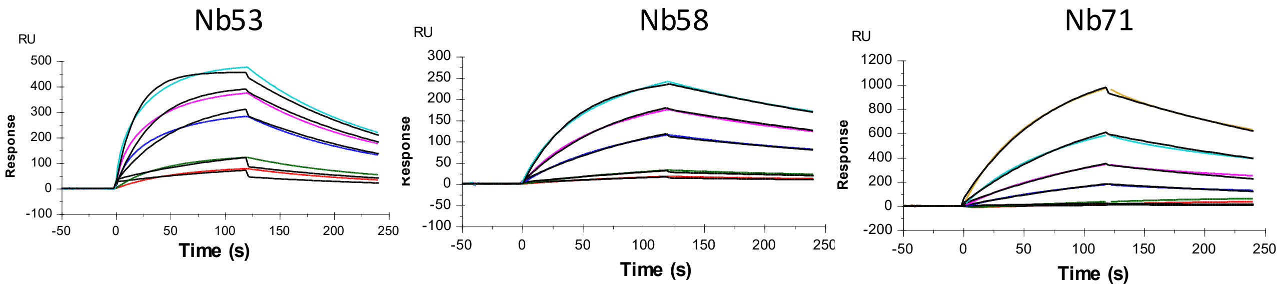


| NB | $k_a$ (1/M.s) | $k_d$ (1/s) | $K_D$ (M) | Rmax (RU) | Capture (RU) | Rmax (%) |
|---|---|---|---|---|---|---|
| **NB53** | 2,31E+05 | 6,10E-03 | **2,64E-08** | 497.0 | 157.11 | 101% |
| **NB58** | 9,94E+04 | 2,79E-03 | **2,81E-08** | 288.7 | 229.35 | 40% |
| **NB71** | 2,55E+04 | 3,43E-03 | **1,35E-07** | 1546 | 988.51 | 50% |

**Supplementary Figure S4. Representative SPR analysis of BasC–nanobody interactions.** Sensorgrams showing the binding of detergent-solubilized BasC to His-tagged Nb53, Nb58 and Nb71 captured on an anti-His-functionalized CM5 sensor chip. BasC was injected at increasing concentrations: 5, 10, 50, 100 and 200 nM for Nb53 and Nb58, and 2.5, 5, 50, 100, 200 and 400 nM for Nb71. Data were fitted using a 1:1 binding model. Association rate constants ($k_a$), dissociation rate constants (k_d), equilibrium dissociation constants (K_D), binding parameters and nanobody capture levels are shown in the table. Rmax (%) represents the fitted maximum response relative to the theoretical maximum response, calculated assuming 1:1 BasC–nanobody binding.

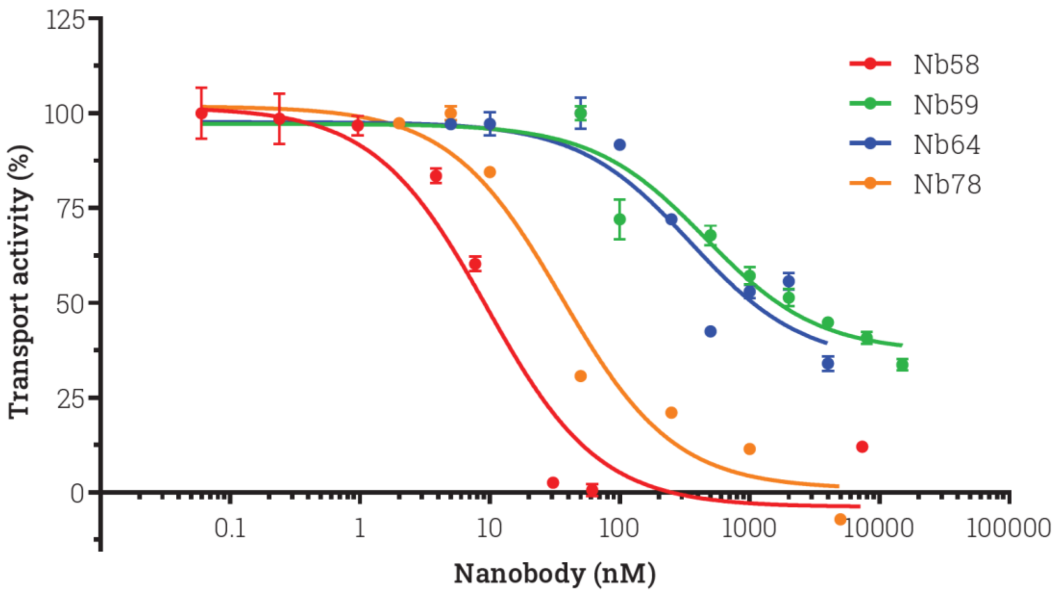


**Supplementary figure S5**. Influx dose responses. Nb58, 59, 64 and 78 influx dose responses. L-[3H]-Ser uptake in 10 µM L-Ser against 4 mM L-Ala filled BasC-GFP-PLs mean ± s.e.m. per each data point from a single experiment triplicates. Lower IC50s were measured for Nb58 and 78 of 9.44 ± 2.42 and 34.8 ± 8.01 nM, respectively. Higher IC50s were measured for Nb59 and 64 of 518 ± 170 and 362 ± 153 nM, respectively.

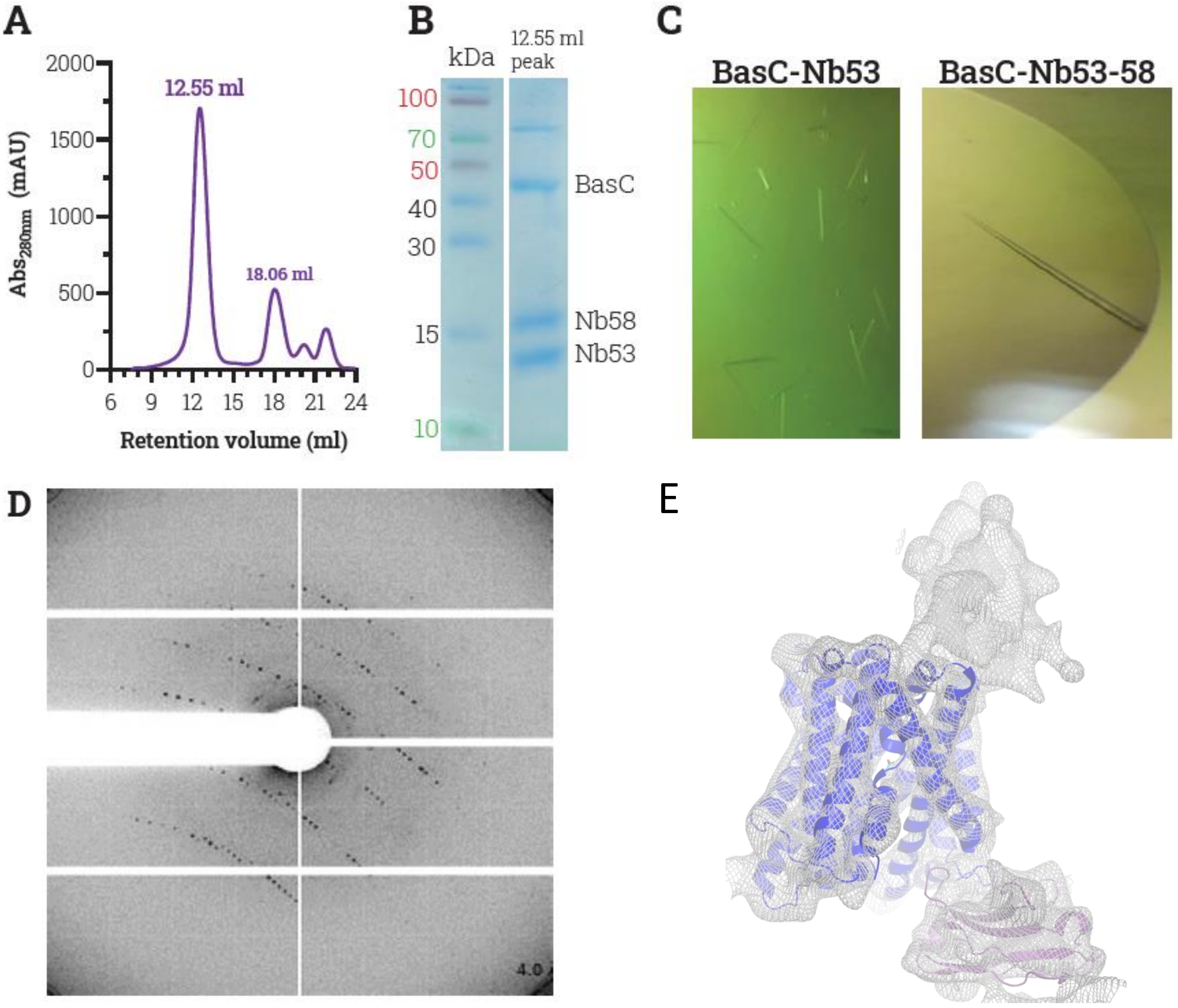


**Supplementary Figure S6. X-ray crystallography of BasC-Nb complexes.**
**(A)** BasC-NB53-58 complex SEC run on a Superdex 200 10/300 GL column in DM buffer. Complex eluted at 12.55 ml, Nb53 and 58 excess, at 18.06 ml. **(B)** SDS-PAGE of BasC-Nb53-58. Both proteins are shown in the 12.55 ml peak lane. Where both Nbs are detected, as BasC. Top band could be BasC dimers. **(C)** Needle-like crystals of BasC-Nb53 (left) and BasC-Nb53-58 crystals (right). Note the improvement on length and thickness. **(D)** Representative diffraction pattern for BasCNb53-58 cristal. **(E)** cryoEM BASC-NB53 model at 2,9 Å fitted in the density of the crystal BASC-NB53-NB58.

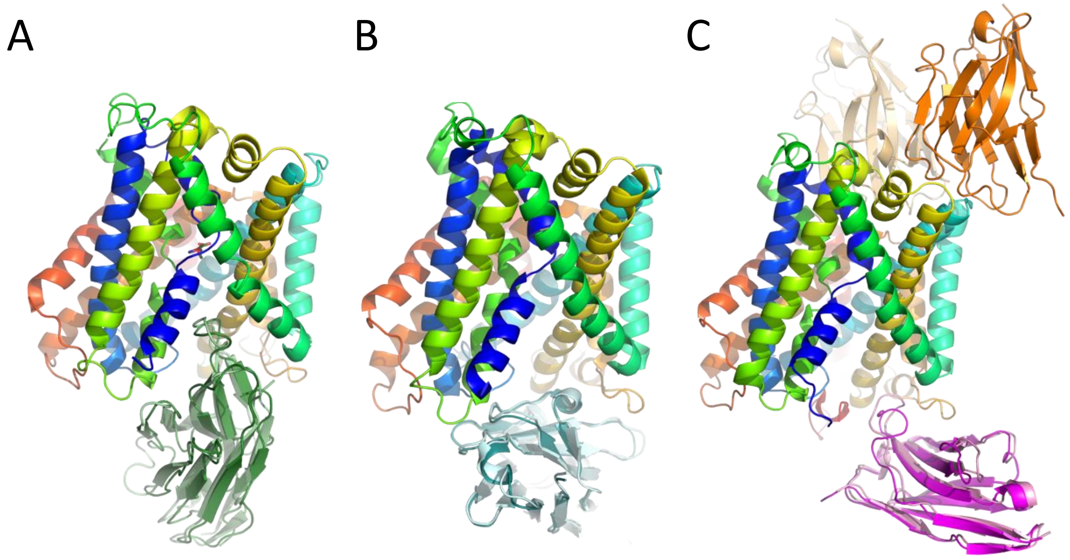


**Supplementary Figure S7. Comparison of AlphaFold3-predicted and experimentally determined BasC–nanobody complexes.** Superpositions of predicted and experimental structures for **(A)** Nb71, **(B)** Nb74 and **(C)** Nb53 and Nb58, aligned using the BasC chains. Only the AlphaFold3-predicted BasC structure is shown for clarity, colored by sequence position. Experimentally determined nanobodies are shown in darker colors, whereas AlphaFold3 predictions are shown in lighter shades. Nb71, Nb74, Nb53 and Nb58 are shown in green, cyan, magenta and orange, respectively. Predictions closely reproduce the experimental binding modes of Nb53, Nb71 and Nb74, whereas the low-confidence Nb58 model differs from its experimentally assigned position in the 6.1 Å crystallographic map. The corresponding ipTM scores are 0.85 for Nb53, 0.16 for Nb58, 0.82 for Nb71 and 0.87 for Nb74. Nb53 and Nb58 were predicted independently and are displayed together for comparison with the experimental ternary complex.
